# Ubiquitin ligase RIPLET mediates polyubiquitination of RIG-I and LGP2 and regulates the innate immune responses to SARS-CoV-2 infection

**DOI:** 10.1101/2021.01.25.428042

**Authors:** Takahisa Kouwaki, Tasuku Nishimura, Guanming Wang, Reiko Nakagawa, Hiroyuki Oshiumi

## Abstract

RIG-I, a cytoplasmic viral RNA sensor, is crucial for innate antiviral immune responses; however, there are controversies about RIG-I’s regulatory mechanism by several ubiquitin ligases and LGP2. Our genetic study revealed that the RIPLET ubiquitin ligase was a general activating factor for RIG-I signaling, whereas another ubiquitin ligase, TRIM25, activated RIG-I in a cell-type-specific manner. These RIPLET and TRIM25 functions were modulated by accessory factors, such as ZCCH3C and NLRP12. Interestingly, we found an additional role of RIPLET in innate immune responses. RIPLET induced delayed polyubiquitination of LGP, resulting in the attenuation of excessive cytokine expression at the late phase. Moreover, RIPLET was involved in the innate immune responses against SARS-CoV-2 infection, a cause of the recent COVID-19 pandemic. Our data indicate that RIPLET fine-tunes innate immune responses via polyubiquitination of RIG-I and LGP2 against virus infection, including SARS-CoV-2.

## Introduction

Cytoplasmic viral RNA sensing initiates innate antiviral immune responses, including type I interferon (IFN) production. RIG-I is a member of the RIG-I-like receptors (RLRs) and senses 5’ tri- and di-phosphate viral double-stranded RNA (dsRNA)^1, 2^. Following viral RNA recognition, RIG-I is associated with the mitochondrial antiviral-signaling protein (MAVS) adaptor protein localized on the outer membrane of mitochondria triggering the expression of cytokines, such as type I IFN and other pro-inflammatory cytokines^3^. Influenza A virus, hepatitis C virus, Sendai virus (SeV), and vesicular stomatitis virus are mainly recognized by RIG-I^4^. *RIG-I* is an IFN-inducible gene expressed in various cell types, such as epithelial cells, dendritic cells, and macrophages^5^. Although RIG-I is crucial for innate antiviral immune responses, excessive RIG-I activation leads to autoimmune disorders^6^. Thus, RIG-I should be tightly controlled by several RIG-I regulators.

The RIG-I protein comprises two caspase activation and recruitment domains (2CARDs), RNA helicase domain, and C-terminal domain (CTD). The protein harbors post-translational modifications, and it has been shown that K63-linked polyubiquitin chains are essential for RIG-I activation^7^. TRIM25 is an E3 ubiquitin ligase that delivers the K63-linked polyubiquitin moiety to RIG-I 2CARDs^7^. The polyubiquitin chains stabilize the 2CARD tetramer structure and promote type I IFN expression^8^. Recent studies have identified several TRIM25 regulators. The ZCCHC3 and NDR2 proteins promote TRIM25-mediated activation of RIG-I, thereby enhancing innate antiviral immune responses^9, 10^. In contrast, NLRP12 inhibits TRIM25 to attenuate RIG-I-mediated signaling^11^.

RIPLET is another ubiquitin ligase that mediates K63-linked polyubiquitination of the CTD and CARDs, and is essential for RIG-I activation^12, 13, 14, 15^. Although many groups have reported TRIM25 as essential for RIG-I activation^1^, several recent studies have challenged this model and shown that RIPLET, but not TRIM25, is necessary for RIG-I-mediated innate immune responses^14, 16, 17^. Besides these contradictions, the ubiquitin ligases MEX3C and TRIM4 have been reported to mediate K63-linked polyubiquitination of RIG-I CARDs^18, 19^. These apparent contradictions are still unresolved^1, 20, 21^.

LGP2 and MDA5 are other RLRs involved in innate antiviral immune responses, including type I IFN expression^22^. LGP2 was once reported as a negative regulator because its over-expression attenuates RIG-I-mediated type I IFN expression^23^. However, LGP2 knockout (KO) exhibited a defect in RIG-I-mediated type I IFN expression after viral infection^24^. LGP2 regulates MDA5 assembly on viral RNA^25^. It has recently been shown that an E3 ubiquitin ligase binds to LGP2^26^; however, the role of LGP2 polyubiquitination in RIG-I activation remains unclear.

Severe acute respiratory syndrome coronavirus 2 (SARS-CoV-2) is a virus causing coronavirus disease 2019 (COVID-19)^27^. The viral spike protein binds to human angiotensin-converting enzyme 2 (ACE2) to infect host cells^27^. The cytoplasmic innate immune response against SARS-CoV-2 remains unclear. Since several viruses have evolved to escape the host innate immune response^1^, it is expected that SARS-CoV-2 can also suppress the innate immune response.

Here, we focused on the role of RIPLET in RIG-I-mediated cytoplasmic antiviral innate immune response. Our genetic data showed that RIPLET is a general activating factor for RIG-I. In contrast, TRIM25 KO exhibited a defect in RIG-I activation in a cell-type-specific manner. We also found RIPLET-mediated polyubiquitination of LGP2. Thus, our data reconcile previous apparent contradictions and indicate that the ubiquitin ligase RIPLET modulates antiviral innate immune responses via the post-translational modification of RIG-I and LGP2.

## Results

### RIPLET KO exhibits severe defects in RIG-I-dependent cytokine expression

Recent studies reported contradictory data related to RIG-I ubiquitination^14, 17^. To address this issue, we generated single-KO, double-KO (DKO), and triple-KO (TKO) cells of TRIM25, RIPLET, MEX3C, and/or TRIM4 using a CRISPR-Cas9 system. We confirmed that CRISPR-Cas9-mediated KO induced several deletion and insertion mutations in each gene and that each KO abolished the expression of TRIM25, MEX3C, TRIM4, and RIPLET (Supplementary Figs. S1a and b). Over-expression of full-length RIG-I leads to its autoactivation, leading to IFN-β promoter activation^12, 13^, and we found that RIPLET KO abolished the full-length RIG-I-mediated IFN-β promoter activation (Fig. 1a). In contrast, neither TRIM25, MEX3C, nor TRIM4 KO failed to reduce the IFN-β promoter activities in HEK293 cells (Fig. 1a).

**Figure 1.**
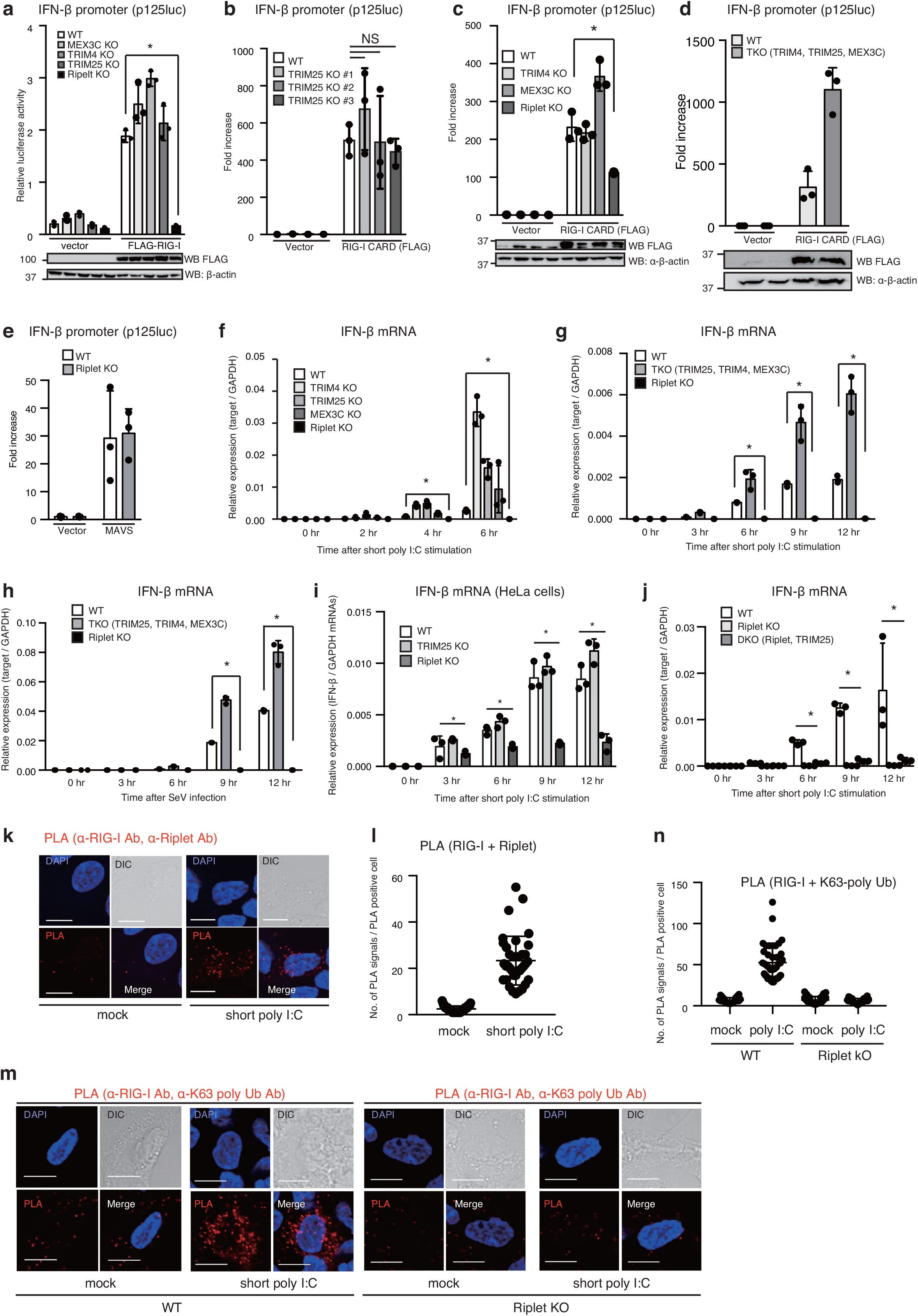
Riplet KO exhibits severe defect in RIG-I-dependent cytokine expression. (a–e) p125luc (IFN-β promoter reporter) plasmid and *Renilla* luciferase vector (internal control) together with indicated expression vectors were transfected into WT and mutant HEK293 cells. The amounts of total DNA in each sample were normalized using an empty vector. After 24 h of transfection, the cells were lysed, and luciferase activities were determined. Western blotting was performed with the indicated antibodies (Abs). The reporter gene activities were measured in three independently isolated TRIM25-KO HEK293 cells (b). The data represent the mean ± SD (n = 3, *p < 0.05, t-test). (f–j) Wild-type and each KO HEK293 (f, g, h, and j) or HeLa (i) cells were transfected with 200 ng/ml of short poly I:C or infected with Sendai virus (SeV) at MOI = 5. Total RNAs were extracted at the indicated time points, and the expression of IFN-β, IP-10, and Ccl5 mRNA was determined by RT-qPCR. The data represent the mean ± SD (n = 3, *p < 0.05, t-test). (k, l) HEK293 cells transfected with 200 ng/ml of short poly I:C for 6 hr were fixed, and PLA was performed with anti-RIG-I and Riplet Abs (k). The number of PLA signals of RIG-I and Riplet were counted in mock- and short poly I:C-stimulated cells (l). (m, n) Wild-type and Riplet-KO HEK293 cells were transfected with 200 ng/ml of short poly I:C for 6 h. Then, the cells were fixed, and PLA was performed with anti-RIG-I and anti-K63 poly-ubiquitin Abs (m). The number of PLA signals were counted in mock- and short poly I:C-stimulated wild-type and Riplet-KO HEK293 cells (n).

As previously reported^7^, TRIM25 over-expression markedly enhanced RIG-I CARDs-mediated type I IFN promoter activation (Supplementary Fig. S1c); however, RIG-I CARDs-mediated promoter activities were not reduced in three independently isolated TRIM25 KO cells (Fig. 1b). We confirmed that endogenous TRIM25 proteins were undetected in the three independent TRIM25 KO clones (Supplementary Fig. S1d). MEX3C KO and TRIM4 KO also failed to suppress promoter activities induced by RIG-I CARDs, whereas RIPLET KO significantly reduced the promoter activities by the RIG-I CARDs (Fig. 1c). TKO of TRIM25, MEX3C, and TRIM4 also failed to mitigate RIG-I CARDs-mediated type I IFN promoter activation (Fig. 1d). We confirmed that RIPLET KO did not reduce the promoter activity driven by MAVS over-expression (Fig. 1e), which weakened the possibility that RIPLET activated the promoter activity in a RIG-I-independent manner.

Next, we investigated cytokine expression in response to RIG-I activation. RIPLET KO abolished the expression of IFN-β, IP-10, and Ccl5 mRNA in response to transfection with short poly I:C, which is a ligand of RIG-I, or Sendai virus (SeV) infection (Figs. 1f-1h). In contrast, KO of each TRIM25, MEXC3C, and TRIM4 failed to reduce the cytokine expression (Fig. 1f and Supplementary Figs. S1e and f). Also, TKO of TRIM25, TRIM4, and MEX3C failed to reduce the cytokine expression in HEK293 cells (Figs. 1g and h, and Supplementary Figs. S1g–j). These data weakened the possibility that TRIM25, MEX3C, and TRIM4 redundantly activated RIG-I to induce type I IFN expression in HEK293 cells. Since some of the TRIM25, MEX3C, TRIM4 KO, and TKO cells exhibited enhanced cytokine expression previously observed by Cadena C et al.^14^, KO of those E3 ubiquitin ligases might lead to suppressor mutations on other gene (s) by an unknown mechanism (see discussion).

Defective RIG-I-mediated type I IFN expression was also detected in RIPLET KO HeLa cells (Fig. 1l). Still, there was no defect in type I IFN expression in TRIM25-KO HeLa cells (Fig. 1i), suggesting that the observation of no defect of RIG-I signaling in TRIM25-KO cells was not specific to HEK293 cells. The DKO of TRIM25 and RIPLET also exhibited a severe defect in short poly I:C-induced expression of IFN-β, IP-10, and Ccl5 mRNA (Fig. 1j and Supplementary Figs. S2a–c).

Next, we observed the subcellular localization of RIPLET, TRIM25, RIG-I, and K63-linked polyubiquitin chains. Proximity ligation assay (PLA) was used to detect the co-localization of two proteins, and the number of endogenous RIG-I and Riplet co-localization PLA signals was increased after stimulation with short poly I:C (Figs. 1k and l), which was consistent with the observation that RIPLET binds to RIG-I after stimulation^12^. Interestingly, several PLA signals were detected on rhodamine-conjugated short poly I:C and mitochondria (Supplementary Figs. S2d and e). Co-localization of RIG-I and K63-linked polyubiquitin chains were also detected by PLA (Fig. 1m). The number of PLA signals was markedly increased after stimulation with short poly I:C, and RIPLET KO abolished the signals (Figs. 1m and n). Several PLA signals were localized on rhodamine-conjugated short poly I:C; however, there were many signals that were not co-localized with the short poly I:C (Supplemental Fig. S2f), implying that ubiquitin-associated RIG-I detached from the ligands after its activation. Our data collectively indicate that the RIPlET, but not TRIM25, MEX3C, nor TRIM4, is essential for RIG-I activation in several cell types, and this notion is consistent with a previous report by Cadena C et al.^14^.

### Cell-type-specific defects of TRIM25 KO cells in RIG-I activation

Although TRIM25 KO unaffected RIG-I-dependent type I IFN expression in HEK293 and HeLa cells, we found that TRIM25 KO reduced IFN-β mRNA expression in A549 cells at the early phase after stimulation with short poly I:C (Fig. 2a), and IP-10 and Ccl5 expression was also reduced by TRIM25 KO at both the early and late phases (Figs. 2b and c). Notably, RIPLET KO also abolished the cytokine expression in response to a RIG-I ligand even in A549 cells (Figs. 2a–c). TRIM25 KO failed to reduce IFN-β mRNA expression after SeV infection (Fig. 2d). In contrast, it moderately reduced IP-10 and Ccl5 mRNA expression in response to SeV infection (Figs. 2e and 2f). To further test the defect of TRIM25 KO in RIG-I activation, we sought to generate additional TRIM25 KO cells and obtained two A549 TRIM25 KO clones (clone #2 and #9). We found that TRIM25 KO significantly reduced the expression of IP-10 and Ccl5 mRNA in two independently isolated TRIM25 KO clones in response to short poly I:C stimulation (Figs. 2g–i), and IP-10 and Ccl5 mRNA expression induced by SeV infection was also reduced in a TRIM25 KO clones (Fig. 2j). Additionally, Ccl5 mRNA expression was reduced by TRIM25 KO in SKOV3 cells after SeV infection (Fig. 2k). PLA signals of endogenous RIG-I and TRIM25 were detected on short poly I:C as was the case with those of RIG-I and Riplet (Fig. 2l). Considering these data, we prefer the interpretation that TRIM25 activates RIG-I in a cell-type-specific manner (see discussion).

**Figure 2.**
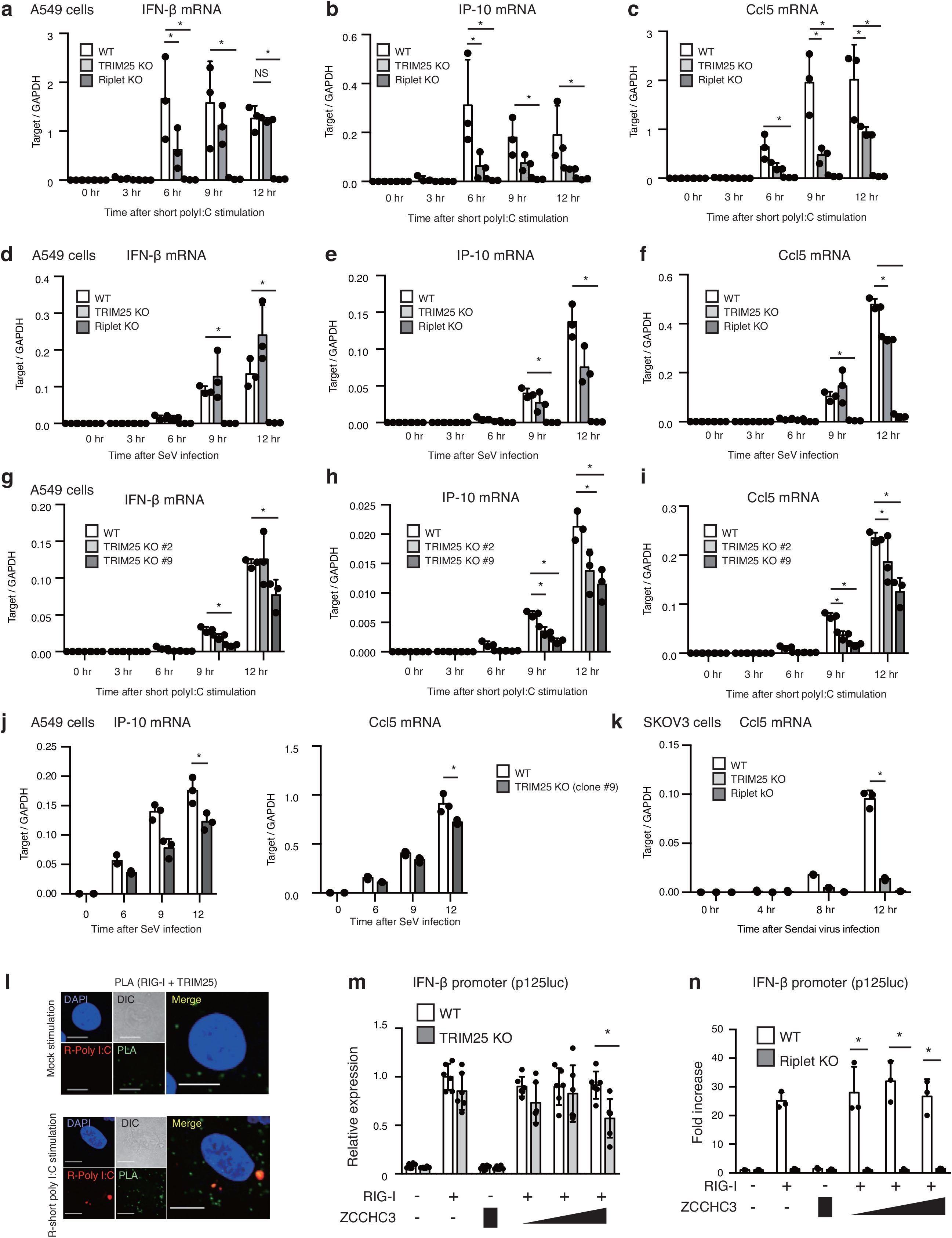
TRIM25 activated RIG-I in a cell-type-specific manner. (a–k) Wild-type and each KO A549 (a–j) or SKOV3 (k) cells were transfected with 200 ng/ml of short poly I:C or infected with SeV at MOI = 5. The expression of cytokine mRNAs were determined by RT-qPCR and normalized to GAPDH. The data represent the mean ± SD (n = 3, *p < 0.05 two-way ANOVA). Three independently isolated TRIM25 KO A549 cells (clone #1: a-f, clone #2 and #9: g–I, clone #9: j) were used in each experiment. (l) A549 cells were transfected with rhodamine-conjugated short poly I:C (R-poly I:C) for 6 h. The cells were then fixed, and PLA (green) was performed with anti-RIG-I and anti-TRIM25 Abs. (m, n) WT, TRIM25 KO, and/or Riplet KO HEK293 cells were transfected with RIG-I and/or ZCCHC3 expressing vectors as indicated. The luciferase activities were determined 24 hr after transfection. The data represent the mean ± SD (*p < 0.05 t-test).

Since recent studies have reported several proteins that regulate TRIM25-mediated RIG-I activation^9, 10, 11^, we hypothesized that the expression of these factors might be required for TRIM25 to activate RIG-I. It has been shown that ZCCHC3 promotes TRIM25-mediated RIG-I activation^10^. ZCCHC3 KO moderately reduced RIG-I-mediated IFN-β and Ccl5 mRNA expression (Supplementary Figs. S2g–i). Although TRIM25 KO did not show any defect in RIG-I signaling in HEK293 cells as described above, TRIM25 KO reduced RIG-I-induced IFN-β promoter activities in ZCCHC3-overexpressing cells (Fig. 2m). We confirmed that RIPLET over-expression enhanced ZCCHC3-mediated RIG-I activation (Supplementary Fig. S2j) and that RIPLET KO abolished RIG-I-mediated type I IFN promoter activation, even in ZCCHC3-overexpressing cells (Fig. 2n). These observations are consistent with the notion that ZCCHC3 promotes TRIM25-mediated RIG-I activation.

NDR2 and NLRP12 have been reported to modulate TRIM25-mediated RIG-I activation^9, 11^. However, NDR2 over-expression augmented RIG-I-mediated IFN-β promoter activation not only in wild-type cells but also in TRIM25 KO cells (Fig. 3a). Also, NLRP12 over-expression reduced RIG-I-mediated signaling in WT and TRIM25 KO cells (Fig. 3b and 3c). These data suggest that NDR2 and NLRP12 regulate RIG-I activities in TRIM25-dependent and -independent manners. Thus, we next investigated the physical interactions between these factors and RIPLET.

**Figure 3.**
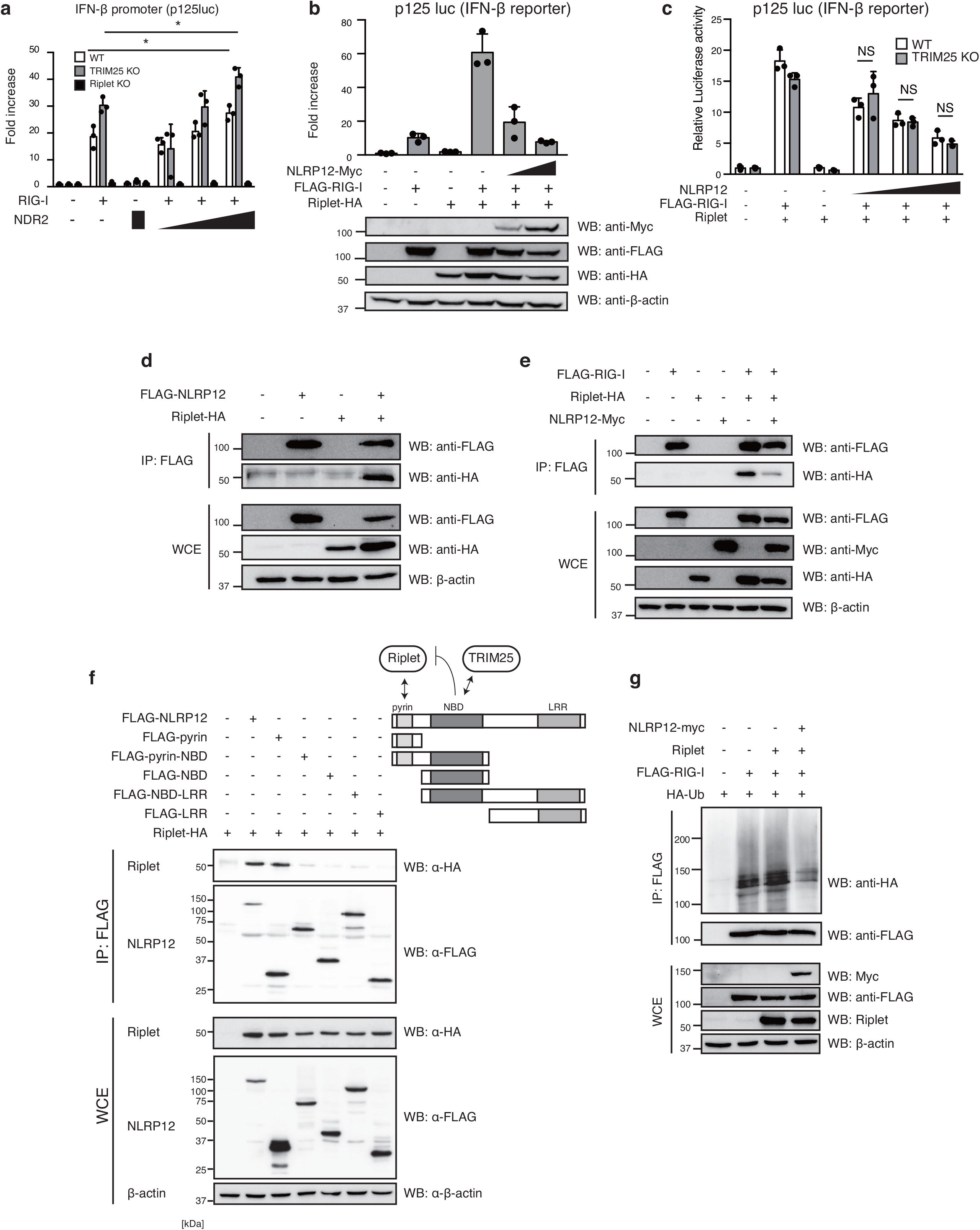
Accessory proteins modulate the Riplet- and TRIM25-medaited RIG-I activation. (a–c) Wild-type, TRIM25 KO, and/or Riplet KO HEK293 cells were transfected with RIG-I, NLRP12, and/or NDR2 expression vectors together with p125luc plasmid and *Renilla* luciferase vector. 24 h after transfection, luciferase activities were determined. The data represent the mean ± SD (n = 3, *p <0.05, t-test). (d) FLAG-tagged NLRP12 and HA-tagged Riplet expression vectors were transfected into HEK293FT cells. WCEs were prepared 24 h after transfection, and immunoprecipitation was performed with anti-FLAG Ab. The proteins were subjected to SDS-PAGE and detected by western blotting with indicated Abs. (e) FLAG-tagged RIG-I, HA-tagged Riplet, and Myc-tagged NLRP12 were transfected into HEK293FT cells. WCEs were prepared 24 h after transfection, and immunoprecipitation was performed with anti-FLAG antibodies. The proteins were subjected to SDS-PAGE and detected by western blotting with the indicated Abs. (f) HA-tagged Riplet and FLAG-tagged NLRP12 fragment expression vectors were transfected into HEK293FT cells. WCEs were prepared 24 h after transfection, and immunoprecipitation was performed with anti-FLAG Ab. The proteins were subjected to SDS-PAGE and detected by western blotting. (g) Myc-tagged NLRP12, Riplet, FLAG-tagged RIG-I, and HA-tagged ubiquitin expression vectors were transfected into HEK293FT cells. WCEs were prepared 24 h after transfection, and immunoprecipitation was performed with anti-FLAG Ab. The proteins were subjected to SDS-PAGE and detected by western blotting with the indicated Abs.

Our immunoprecipitation assay showed that RIPLET was co-immunoprecipitated with NLRP12 (Fig. 3d), and NLRP12 expression reduced the physical interaction between RIG-I and RIPLET (Fig. 3e). TRIM25 binds to the nucleotide-binding domain (NBD) of NLRP12^11^. Our immunoprecipitation assay with truncated forms of NLRP12 showed that RIPLET bound to the pyrin domain of NLRP12, and the NBD inhibited the interaction (Fig. 3f). Moreover, NLRP12 expression inhibited RIPLET-mediated RIG-I polyubiquitination of RIG-I (Fig. 3g). These data suggest that the accessory factors are involved in both RIPLET- and TRIM25-mediated RIG-I activations.

### RIPLET mediates K63-linked polyubiquitination of LGP2 C-terminal region

Next, we focused on the role of LGP2, because unknown function of LGP2 may underlie the controversy. LGP2 is a member of the RLRs. Its over-expression reduced RIG-I-mediated IFN-β promoter activation but enhanced MDA5-mediated IFN-β Promoter activation (Supplementary Figs. S3a and b). However, LGP2 KO significantly reduced mRNA expression of IFN-β and IP-10 in response to short poly I:C and SeV infection (Figs. 4a–d). Since there are similarities between the CTDs of RIG-I and LGP2^28^, we investigated the physical interaction between RIPLET and LGP2 and found that the RIPLET protein was co-immunoprecipitated with LGP2 (Figs 4e and f). Additionally, PLA signals of RIPLET-HA and FLAG-LGP2 were detected (Fig. 4g). Interestingly, LGP2 harbored polyubiquitin modifications, and RIPLET enhanced LGP2 polyubiquitination (Fig. 4h). To investigate whether the modification was K63-linked polyubiquitination, K63-only ubiquitin, in which all K residues of ubiquitin except K at 63 were replaced with R, was utilized to detect LGP2 ubiquitination. K63-only ubiquitin was efficiently incorporated into LGP2, and that the polyubiquitin bands were increased by RIPLET expression (Fig. 4i). Conversely, RIPLET KO reduced the polyubiquitination of LGP2 (Fig. 4j). The PLA signals of endogenous LGP2 and K63-polyubiquitin were observed. The number of signals was increased by stimulation with short poly I:C, and RIPLET KO abolished the PLA signals (Fig. 4k). Additionally, the signals were detected on transfected rhodamine-short poly I:C (Fig. 4l). These data suggested that RIPLET mediated K63-linked polyubiquitination of LGP2.

**Figure 4.**
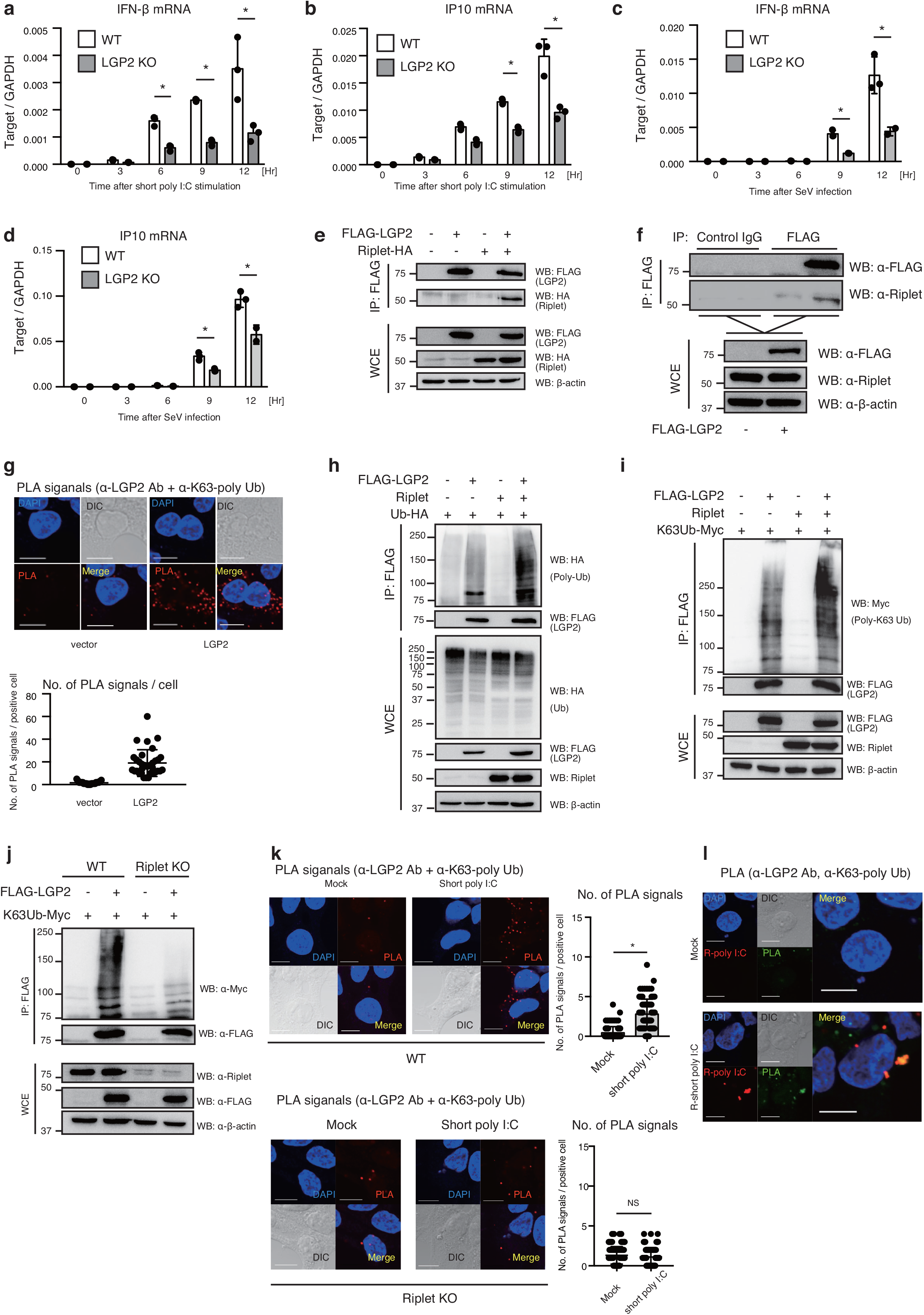
K63-linked polyubiquitination of LGP2. (a–d) Wild-type and LGP2-KO HEK293 cells were transfected with 200 ng/ml of short poly I:C or infected with Sendai virus (SeV). Total RNAs were extracted as the indicated time points, and IFN-β and IP-10 mRNA expression was determined by RT-qPCR. The data represent the mean ± SD (n = 3, *p < 0.05, t-test). (e) HEK293FT cells were transfected with FLAG-tagged LGP2 and HA-tagged Riplet. WCEs were prepared 24 h after transfection, and immunoprecipitation was performed with anti-FLAG Ab. The proteins were subjected to SDS-PAGE and detected by western blotting as indicated. (f) HEK293FT cells were transfected with FLAG-tagged LGP2. WCEs were prepared 24 h after transfection, and immunoprecipitation was performed with anti-FLAG Ab or control IgG. The proteins were subjected to SDS-PAGE and detected with the indicated Abs. (g) HEK293 cells were transfected with an empty and LGP2 expression vector. The cells were fixed after 24 h of transfection, and PLA was performed with anti-LGP2 and anti-K63 polyubiquitin Abs (upper panel). The number of PLA signals was counted (lower panel). (h) FLAG-tagged LGP2, Riplet, and HA-tagged ubiquitin expression vectors were transfected into HEK293FT cells. WCEs were prepared 24 h after transfection, and immunoprecipitation was performed with anti-FLAG Ab. The proteins were subjected to SDS-PAGE and detected by western blotting with the indicated Abs. (i) FLAG-tagged LGP2, Riplet, and Myc-tagged K63 only ubiquitin, in which all K amino acids except K63 were replaced with R amino acid, were transfected into HEK293FT cells. WCEs were prepared 24 h after transfection, and immunoprecipitation was performed with anti-FLAG Ab. The proteins were subjected to SDS-PAGE and detected by western blotting, as indicated. (j) FLAG-tagged LGP2 and Myc-tagged K63 only ubiquitin were transfected into HEK293 Wild-type and Riplet-KO cells. WCEs were prepared 24 h after transfection, and immunoprecipitation was performed with anti-FLAG Ab. The proteins were subjected to SDS-PAGE and detected by western blotting, as indicated. (k) Wild-type and Riplet-KO HEK293 cells were stimulated with 200 ng/ml of short poly I:C for 6 h. PLA was performed with anti-LGP2 and anti-K63 polyubiquitin Abs. The number of the PLA signals was counted. (l) HEK293 cells were stimulated with mock or rhodamine-conjugated short poly I:C (R-poly I:C) for 6 h. PLA (green) was performed with anti-LGP2 and anti-K63 polyubiquitin Abs. The PLA signals are shown in green.

Since the immunoprecipitation assay and PLA suggested that RIPLET ubiquitinated LGP2, we performed a mass spectrometry analysis to determine the ubiquitination sites of LGP2. We found that K39, K532, K599, and K629 of LGP2 were ubiquitinated in RIPLET-expressing cells (Fig. 5a and Supplementary Fig. S3c). The alignment of LGP2 and RIG-I showed that K532 and K599 of LGP2 were comparable with K788 and K851 of RIG-I, which are ubiquitinated by RIPLET (Supplementary Fig. S3d). Immunoprecipitation assay showed that RIPLET preferentially bound to the helicase domain of LGP2, although the CTD is also associated with RIPLET (Fig. 5b), which is consistent with a recent study showing that ubiquitin ligases bind to the helicase domain of RLRs^26^. The 3KR amino acid substitutions within the CTD reduced the polyubiquitination of LGP2 CTD (Fig. 5c), but residual polyubiquitination was still detected.

**Figure 5.**
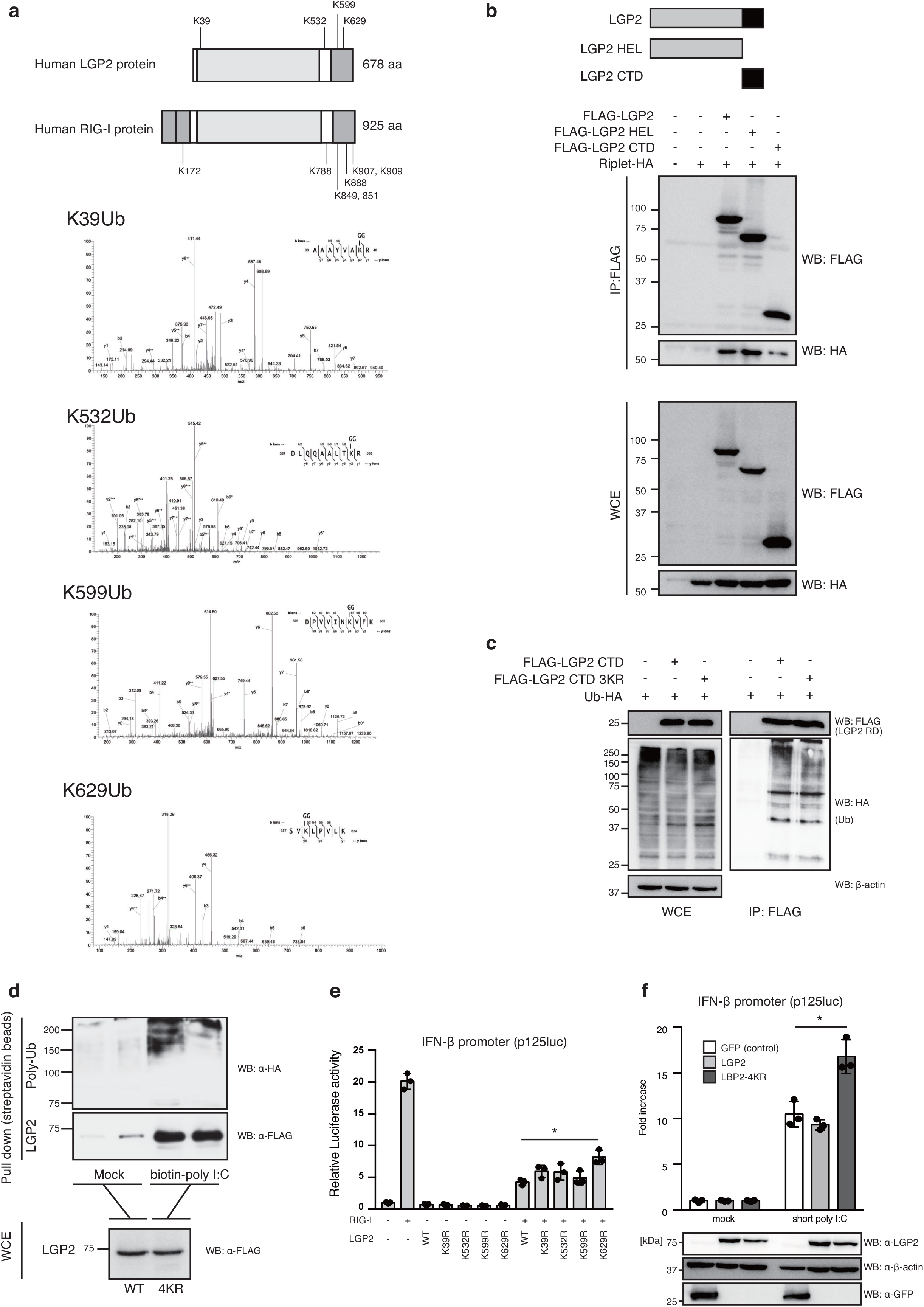
Ubiquitination sites of LGP2. (a) Liquid chromatography-tandem mass spectrometry (LC/LC-MS/MS) of LGP2 in Riplet-expressing cells. Schematic representation of ubiquitination sites of RIG-I and LGP2 were shown in upper panel. (b) FLAG-tagged LGP2 fragments and HA-tagged Riplet expression vectors were transfected into HEK293FT cells. WCEs were prepared 24 h after transfection, and immunoprecipitation was performed with anti-FLAG Ab. The proteins were subjected to SDS-PAGE and detected by western blotting, as indicated Abs. (c) HA-tagged ubiquitin, FLAG-tagged LGP2 CTD, and FLAG-tagged LGP2 CTD 3KR expression vectors were transfected into HEK293FT cells. WCE were prepared 24 h after transfection, and immunoprecipitation was performed with anti-FLAG Ab. The proteins were subjected to SDS-PAGE and detected by western blotting, as indicated Abs. (e) HA-tagged ubiquitin and FLAG-tagged wild-type and 4KR LPG2 expression vectors were transfected into HEK293FT cells. WCEs were prepared, and a pull-down assay with mock or biotin-conjugated poly I:C was performed. The proteins were subjected to SDS-PAGE and detected by western blotting, as indicated Abs. (f) RIG-I and/or wild-type and KR mutants of LGP2 were transfected into HEK293 cells together with p125luc plasmid and *Renilla* luciferase vector. Luciferase activities were determined 24 h after transfection. The data represent the mean ± SD (n = 3). (g) HE293 cells stably expressing GFP (control), LGP2, LGP2-4KR were generated using lentivirus vectors. The cells were transfected with p125luc plasmid and *Renilla* luciferase vectors and then stimulated with short poly I:C. WCEs were prepared 24 h after transfection and stimulation, and luciferase activities were determined. The proteins were subjected to SDS-PAGE and detected by western blotting with the indicated Abs. The data represent the mean ± SD (n = 3, *p < 0.05, t-test).

To investigate the physiological significance of LGP2 polyubiquitination, we constructed a LGP2-4KR mutant in which K39, K532, K599, and K629 of LGP2 were replaced with the R residue. Both non-ubiquitinated and polyubiquitinated wild-type and LGP2-4KR mutant proteins were recovered by pull-down assay with poly I:C-conjugated streptavidin beads (Fig. 5d), suggesting that polyubiquitination did not affect the RNA-binding activity of LGP2. To investigate the effect of LGP2 ubiquitination on cytokine expression, we constructed LGP2 KR mutants. We observed a moderate increase in RIG-I-mediated IFN-β promoter activation with the K39R or K532R substitution, and the K629R mutation significantly increased the promoter activities (Fig. 5e). We generated cells that stably expressed wild-type LGP2 or LGP2-4KR using a lentivirus vector^29^. Stable expression of wild-type or mutant LGP2 unaffected the IFN-β promoter activities in non-stimulated cells (Fig. 5f). Interestingly, the LGP2-4KR mutant protein induced the promoter activity more than the wild-type protein in response to a RIG-I ligand (Fig. 5f). These data suggested that LGP2 ubiquitination negatively affects type I IFN expression. Next, we determined the mRNA expression of cytokines in response to stimulation with a RIG-I ligand and SeV infection. The 4KR mutation augmented the cytokine expression at late time points but not at the early time points (Figs. 6a–f). Time course analyses showed that the PLA signals of LGP2 and K63-polyubiquitin increased more slowly than RIG-I and K63-polyubiquitin (Figs. 6g–i). Influenza A virus and SARS-CoV-2 are suspected of inducing cytokine storm, in which excessive cytokine production is harmful to the host^30, 31^. RIPLET-mediated K63-linked polyubiquitination of LGP2 is expected to attenuate excessive cytokine expression at the late phase.

**Figure 6.**
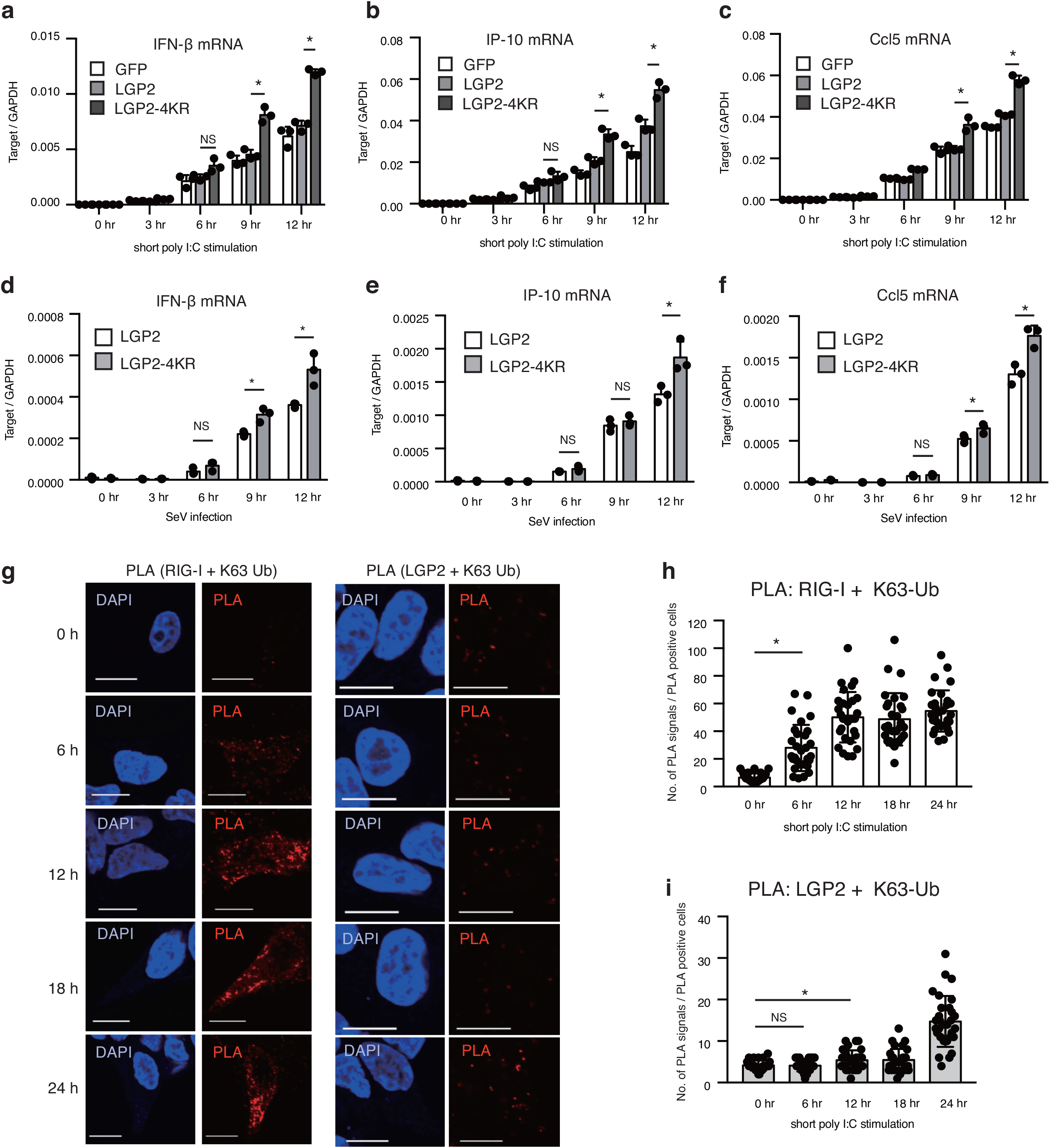
Ubiquitination of LGP2 attenuates the cytokine expression at a late phase. (a–f) HEK293 cells stably expressing GFP, LGP2, and LGP2-4KR were transfected with 200 ng/ml of short poly I:C or infected with Sendai virus (SeV) at MOI = 5. Total RNAs were extracted at the indicated time points. The expression of IFN-β mRNA was determined by RT-qPCR. The data represent the mean ± SD (n = 3, *p < 0.05, t-test). (g–i) HEK293 cells were stimulated with 200 ng/ml of short poly I:C. The cells were fixed at the indicated time points, and PLA was performed with anti-RIG-I, anti-LGP2, and/or anti-K63 polyubiquitin Abs (J). The PLA signals of RIG-I/K63-Ub (H) and LGP2-K63/Ub (I) were counted.

### RIPLET plays a crucial role in innate immune responses against SARS-CoV-2

Studies have shown that RIPLET plays a crucial role in the innate immune response to viral infection, such as influenza A, vesicular stomatitis virus, Sendai, and hepatitis C virus^12, 32^. Since SARS-CoV-2 is a virus causing the COVID-19 pandemic, we investigated whether RIPLET is involved in the innate immune response to SARS-CoV-2. First, we investigated whether RIG-I recognizes SARS-CoV-2 RNA. VeroE6/TMPRSS2 cells were infected with SARS-CoV-2, and the total RNA of infected cells was extracted. Extracted RNAs from infected and non-infected cells were transfected into HEK293 cells, and cytokine expression was measured. Total RNAs from virus-infected cells but not those from uninfected cells induced cytokine expression (Fig. 7a and Supplementary Figs. S4a and b), suggesting that RNAs of SARS-CoV-2-infected cells can induce the innate immune response. Interestingly, either RIG-I KO or MDA5 KO markedly reduced cytokine expression in response to viral infected cell RNAs (Fig. 7b). The dependency on both RIG-I and MDA5 has been reported in the response to Japanese encephalomyelitis virus infection^33^. Our data are consistent with a recent report showing that MDA5 plays a crucial role in the innate immune response to SARS-CoV-2.^34^.

**Figure 7.**
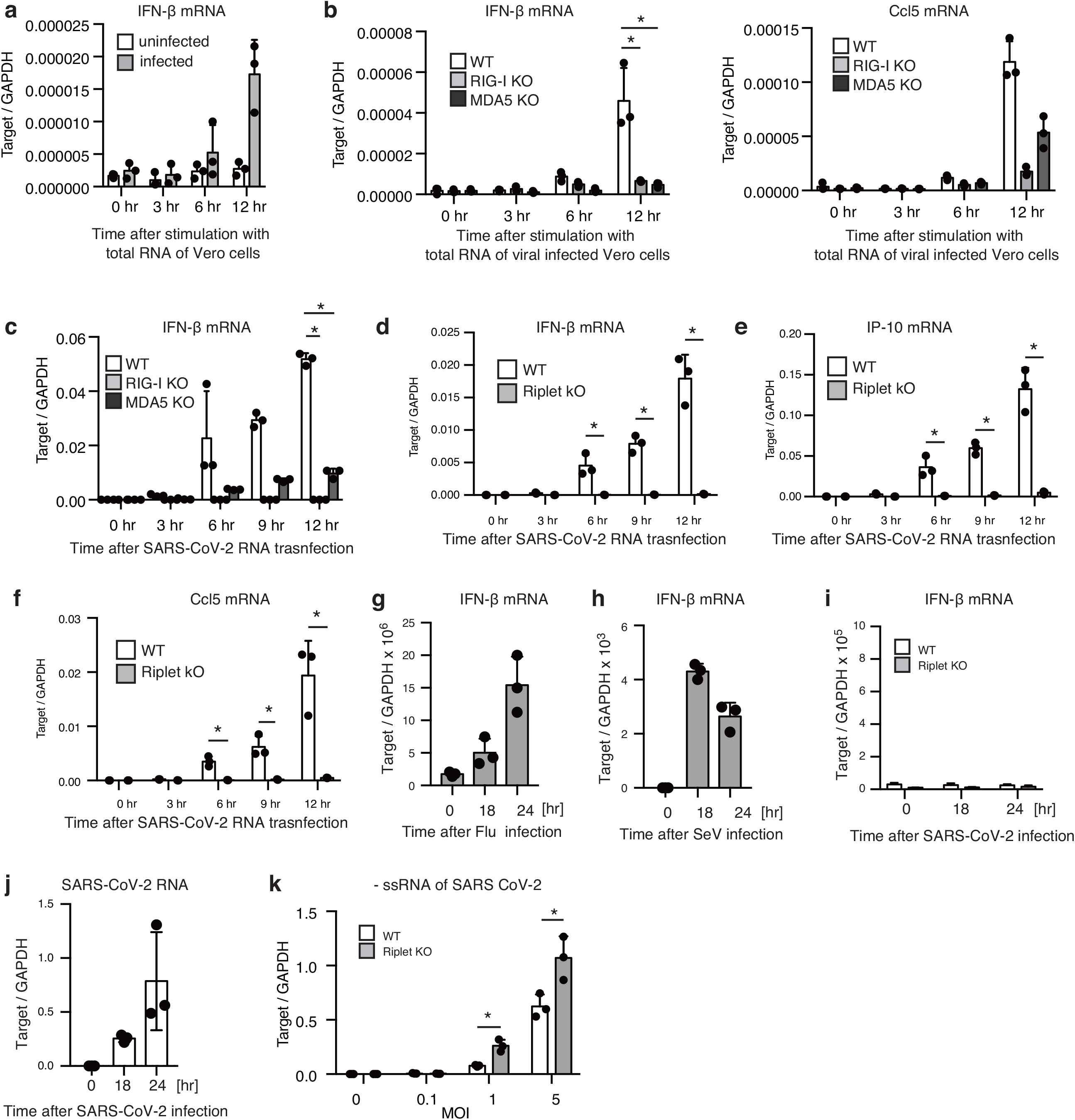
Riplet plays a crucial role in antiviral response against SARS-CoV-2. (a) Total RNAs were extracted from VeroE6/TMPRSS2 cells infected with or without SARS-CoV-2 for 24 hr. 1 µg of extracted RNA was transfected into HEK293 cells, and the cytokine expression at indicated time points were determined by RT-qPCR (n = 3). (b) 1 µg of Total RNA from VeroE6/TMPR22S infected with SARS-CoV-2 were transfected into WT, RIG-I KO, and MDA5 KO cells. The cytokine expression at indicated time points were determined by RT-qPCR (n = 3, *p<0.05, t-test). (c–f) The viral RNA segment (24001–25000) was transfected into WT, RIG-I KO, MDA5 KO, and Riplet KO cells. Total RNA was extracted at indicated time points, and the expression of the cytokines were determined by RT-qPCR. The data represents the mean ± SD (n = 3, *p < 0.05, t-test). (g–j) HEK293 were infected with influenza A virus (g), SeV (h), and SARS-CoV-2 (i, j) at MOI = 1. Total RNA was extracted at indicated time points, and the cytokine mRNA and viral RNA were determined by RT-qPCR. The data represents the mean ± SD (n = 3). (k) HEK293 cells stably expressing ACE2 were infected with SARS-CoV-2 at indicated MOI for 18 hr. IFN-β mRNA, IP-10 mRNA, and negative stranded viral RNA of SARS-CoV-2 were determined by RT-qPCR. The data represents the mean ± SD (n = 3, *p<0.05, t-test).

Second, we investigated which viral RNA genome segments are recognized by the cytoplasmic innate immune system. Contiguous genomic viral RNA segments of SARS-CoV-2 were prepared using T7 RNA polymerase shown in Supplemental Fig. S4c. Each viral RNA segment was transfected into HEK293 cells, and IFN-β mRNA expression was measured. Interestingly, two viral RNA segments, 24001–2500 and 27001–28000, significantly induced the type I IFN expression. We found that either RIG-I KO or MDA5 KO was enough to reduce the cytokine expression to the viral RNA segment of SARS-CoV-2 (Fig. 7c and Supplementary Figd S4d). Interestingly, RIPLET KO abolished the cytokine expression in response to the SARS-CoV-2 RNA segment (Figs. 7d–f), suggesting the crucial role of RIPLET in the innate immune response during SARS-CoV-2 infection.

Third, we investigated the cytokine expression after viral infection. Although influenza A virus or SeV infection at MOI = 1 induced IFN-β mRNA expression, SARS-CoV-2 infection failed to induce the cytokine expression (Fig. 7g–i and Supplementary Fig. S4e). We confirmed that SARS-CoV-2 replicated in HEK293 cells (Fig. 7j). Recent studies have shown that viral proteins of SARS-CoV-2 can suppress the innate immune responses, and we also found that several viral proteins suppressed the RIG-I-mediated IFN-β promoter activation (Supplementary Fig. S4f). To further address the role of RIPLET in the innate immune response, we generated human ACE2 overexpressing HEK293 cells (HEK293-ACE2) to increase the infectivity of SARS-CoV-2 to HEK293 cells as previously reported^27, 35^. In HEK293-ACE2 cells, SARS-CoV-2 infection induced moderate expression of type I IFN expression, and RIPLET KO reduced the cytokine expression (Supplementary Fig. S4g). Interestingly, RIPLET KO significantly increased the cytoplasmic viral levels after SARS-CoV-2 infection (Fig. 7k). Our data collectively indicate that RIPLET is required for the antiviral innate immune response against SARS-CoV-2.

## Discussion

RIG-I is a cytoplasmic viral RNA sensor that is essential for innate antiviral immune responses. Although there is controversy about RIG-I regulators, our data revealed that RIPLET ubiquitin ligase is a general factor for RIG-I activation, and TRIM25 KO exhibited a defect in RIG-I activation in a cell-type-specific manner. Moreover, several accessory factors affected not only TRIM25-but also RIPLET-mediated RIG-I activation. We also found that RIPLET induced delayed polyubiquitination of LGP2, which attenuates the excessive cytokine expression. Furthermore, RIPLET was essential for the antiviral innate immune response against SARS-CoV-2, which is a cause of the recent COVID-19 pandemic. These data elucidated a crucial role of the RIPLET ubiquitin ligase in regulating antiviral innate immune responses.

K63-linked polyubiquitination of RIG-I is an essential step to trigger the expression of type I IFN and other pro-inflammatory cytokines. Many studies have reported that TRIM25 delivers the K63-linked polyubiquitin moiety to RIG-I, thereby inducing type I IFN expression in vitro and in vivo^1^. However, recent studies have challenged this model^14, 16, 17^. Here, we carefully assessed the ubiquitin ligases’ role using single-, double-, and triple-KO cells and the obtained data that reconciled the apparent discrepancy. As reported by Cadena et al., we found that RIPLET KO severely reduced RIG-I-dependent cytokine expression in any cell type as far as we tested. In contrast, TRIM25 KO attenuated the RIG-I-dependent cytokine expression only in a cell-type-specific manner. Considering that TRIM25 has been considered essential for RIG-I activation in any cell types^7^, our finding would reconcile previous apparent discrepancy of why TRIM25 KO exhibited a defect in RIG-I activation in some studies but did not in other studies^1, 21^.

In addition to RIPLET and TRIM25, MEX3C and TRIM4 were reported to be involved in RIG-I ubiquitination. Unlike TRIM25 and RIPLET, we could not see any effect of MEX3C KO or TRIM4 KO on RIG-I activation at least in our experimental conditions. We do not exclude the possibility that MEX3C and TRIM4 might be required for RIG-I activation in a cell-type-specific manner, as seen with TRIM25. Recent studies have reported several factors that regulate TRIM25 functions. Since TRIM25 decreased RIG-I-mediated type I IFN expression in HEK293 cells expressing ZCCHC3; it is expected that some types of immortalized cells lack the factors’ expression required for TRIM25-mediated RIG-I activation.

Some of TRIM25 KO, MEX3C KO, and/or TRIM4 KO exhibited enhanced cytokine expression in response to RIG-I stimulation in our experimental conditions. These observations were also observed in a previous study by Cadena et al^14^. Thus, it is suspected that some suppressor mutations occurred on other unknown genes during the isolation of those KO cells. Considering that ubiquitin ligases target several proteins, it is not surprising that suppressor mutations occur in ubiquitin ligase KO cells^36, 37^. Those spontaneous suppressor mutations might affect the phenotypes of TRIM25 KO cells. Thus, we prefer the interpretation that TRIM25 activates RIG-I in a cell-type-specific manner rather than the model that TRIM25 is not involved in RIG-I activation. Further studies using TRIM25 KO mice and TRIM25 KO primary cells are required to settle the dispute.

The decrease in of IP-10 and Ccl5 expression was more significant than that of IFN-β in TRIM25 KO cells. The MAVS adaptor induces cytokine expression via mitochondria or peroxisome^38^. Peroxisomal MAVS preferentially activates the transcription of interferon-stimulated genes including IP-10, using IRF1 and IRF3, whereas mitochondrial MAVS activates only IRF3, thereby inducing type I IFN expression^38^. Considering that TRIM25 KO exhibited more severe defects in IP-10 and Ccl5 expression than that of type I IFN, it is possible that TRIM25-mediated polyubiquitin chain preferentially activates peroxisomal MAVS. Since RIPLET functions as a co-receptor for RIG-I and an E3 ubiquitin ligase for RIG-I, it is still possible that K63-linked polyubiquitin chains are important for RIG-I-induced IP-10 and Ccl5 expression, and the co-receptor function of RIPLET is required for the type I IFN expression. This is also supported by the notion that the mutations on the ubiquitinated RIG-I residues exhibited a relatively moderate phenotype compared with RIPLET KO. Further studies are required to address this issue.

Abnormal activation of RIG-I causes Singleton-Merten syndrome, which is a multisystem autoimmune disorder, and an excessive innate immune response is harmful to the host^6^. We identified an additional role of RIPLET: it mediated K63-linked polyubiquitination of LGP2, leading to the attenuation of excessive type I IFN expression at the late phase. LGP2 was once reported as a negative regulator^22, 23^, but later studies have shown that LGP2 plays a positive role in RIG-I-mediated cytokine expression^24^. This study showed that LGP2 KO reduced RIG-I-mediated cytokine expression, but inhibition of LGP2 ubiquitination enhanced cytokine expression. These observations indicate that LGP2 has a dual role in the regulation of RIG-I signaling. This model can explain why ectopically expressed LGP2 exhibits a negative effect because LGP2 over-expression led to LGP2 polyubiquitination that attenuated RIG-I signaling. K63-linked polyubiquitination of LGP2 was detected only at the late phase after stimulation of RIG-I Thus, delayed polyubiquitination of LGP2 is expected to be a negative feedback mechanism to suppress excessive cytokine expression harmful to the host.

Cytokine storm is suspected to be a cause of mortality in patients with COVID-19^30^. Indeed, treatment with tocilizumab, a monoclonal antibody against IL-6, reduces the mortality of patients with COVID-19^39^. Therefore, we prefer the interpretation that both RIPLET-dependent RIG-I and LGP2 ubiquitination would be essential to protect the host from viral infection. Since SARS-CoV-2 infection hardly induced type I IFN and pro-inflammatory cytokine expression in HEK293 and HEK293-ACE2 cells, RIPLET-dependent cytoplasmic innate immune system seems to be uninvolved in harmful excessive pro-inflammatory cytokine production in patients with COVID-19. Toll-like receptors, such as TLR3, TLR7, and TLR8, might be involved in excessive pro-inflammatory cytokine production. We do not exclude the possibility that some kinds of cells produce pro-inflammatory cytokine via RIPLET-dependent cytoplasmic innate immune response during SARS-CoV-2 infection. However, our observation that several viral proteins can suppress the cytoplasmic innate immune response weakens this possibility.

Although there are several contradictory reports related to RIG-I ubiquitination and the role of LGP2 in innate immune response, our data could reconcile those apparent discrepancies as discussed above. The molecular mechanism of RIPLET-dependent regulation of RIG-I and LGP2 would be required to fully understand the intracellular innate immune responses against viral infections including SARS-CoV-2.

## Acknowledgments

We thank our laboratory members for helpful discussion. This work was supported in part by Grants-in-Aid from the Ministry of Education, Culture, Sports, Science, and Technology (MEXT) and Japan Agency for Medical Research and Development (AMED).

## Data availability

Our original data have been deposited to Mendeley Data (https://data.mendeley.com/datasets/nhbj996m4h/draft). Further information and requests for resources, data, and reagents should be directed to lead contact Hiroyuki Oshiumi.

## Author contributions

HO and TK designed the experiments. RN performed mass spectrometry analysis. TK, TN, and GW performed other experiments. HO and TK wrote the manuscript.

## Materials and Methods

### Cell Lines

TRIM4-, TRIM25-, MEX3C-, Riplet-, ZCCHC3-, and LGP2-KO cells were generated using a CRISPR/Cas9 system. The guide sequence (See Table S1) was cloned into a BbsI restriction site of a pX459 plasmid carrying Cas9 and puromycin resistance genes. The plasmids carrying the guide sequence and guide sequence were transfected into HEK293, A549, and HeLa cells. After 36 h of transfection, the cells were treated with 1 µg/mL of puromycin for 3 days, and then were cultured in fresh medium without puromycin for 2 days. Finally, cells were seeded into 96-well plates and cultured for 7–10 days to obtain a single clone, which was passed into 24-well plates. Western blotting was performed to confirm KO of the target gene. To obtain the TKO (TRIM4 KO, TRIM25 KO and MEX3C KO) HEK293 cell line, TRIM4-KO cells were transfected with pX459 plasmids carrying TRIM25 guide RNA and single clones were obtained, as described above. Finally, TRIM4 and TRIM25 KO HEK293 cells were transfected with pX459 plasmids carrying with the MEX3C guide RNA, and single clones were established using the abovementioned method.

### Constructs

cDNAs encoding full-length ORFs of ZCCHC3, NDR2 and LGP2 were amplified with cDNA library from HEK293 cells, and were cloned into the *Kpn*I restriction site of pEF-BOS using an In-Fusion Cloning Kit (TaKaRa Bio.). A FLAG-tag sequence was inserted just after the start codon. A cDNA of NLRP12 was amplified with a cDNA library derived from THP1 cells and was cloned into the *Kpn*I restriction site of pEF-BOS using an In-Fusion Cloning Kit. A FLAG-tag sequence was inserted just after the start codon. To obtain a Myc-tagged NLRP12 expression vector, the ORF of NLRP12 was amplified from the pEF-BOS-FLAG-NLRP12 vector and cloned into the *Bam*HI and *Xho*I restriction site of pCDNA3 using an In-Fusion Cloning Kit. A Myc-tag sequence was inserted into just before the stop codon. Truncation mutants of NLRP12 and LGP2 were amplified with abovementhoned constructed plasmids and cloned into the *Kpn*I restriction site of pEF-BOS using an In-Fusion Cloning Kit. A FLAG-tag sequence was inserted just after the start codon. The KR mutations were introduced into the LGP2 gene on the expression plasmid using overlapped forward and reverse primers. Full-length LGP2 KR mutants were amplified using two fragments as templates and cloned into the pEF-BOS-FLAG vector.

### Cell cultures and viruses

HEK293 cells were maintained in DMEM (low Glc) supplemented with 10% heat-inactivated FCS and penicillin-streptomycin solution. HEK293FT and A549 cells were incubated in DMEM (high Glc) supplemented with 10% heat-inactivated FCS and penicillin-streptomycin solution. HeLa cells were cultured in MEM supplemented with 10% heat-inactivated FCS and penicillin-streptomycin solution. These cells were grown in a humidified incubator maintained at an atmosphere 5 % CO_2_. SeV was amplified using Vero cells, and the number of plaque-forming units was determined by a plaque assay. The cells were infected with SeV at a multiplicity of infection of 5 to determine the cytokine expression after SeV infection. SARS-CoV-2 was kindly provided from national institute of infectious diseases in Japan ^40^. SARS-CoV-2 was amplified with VeroE6/TMPRSS2 cells. Viral titers were determined by TCID50 assay according to Behrens-Karber’s method. Total RNA of VeroE6/TMPRSS2 cells infected with SARS-CoV-2 was isolated with TRIZOL, and cDNA was prepared using random primer. The cDNA fragments encoding the viral RNA segments were amplified with the primer described in supplemental Table S1. The viral RNA segments were synthesized with T7 RNA polymerase. The viral RNA segment (5001–6000) was not able to be amplified by unknown reason. The experiments using SARS-CoV-2 were performed in BSL3 level facilities according to approved protocols in Kumamoto University.

### Western blot analysis

Cells were washed with PBS and lysed with NP-40 lysis buffer (20 mM tris-HCl [pH7.5], 125 mM NaCl, 1 mM EDTA, 10% Glycerol, 1% NP-40, 30 mM NaF, and 5 mM Na_3_VO_4_) in the presence of complete Protease Inhibitor Cocktail (Roche). The cell lysates were incubated on ice for 30 min and were then centrifuged for 20 min at 15,000 rpm at 4°C. The supernatants were transferred to 1.5 ml tubes and suspended in 2 x Laemmil sample buffer containing β-mercaptoethanol. The cell lysates were boiled for 5 min at 95°C. All samples and a protein marker (Bio-Rad) were each loaded into separate wells for SDS-PAGE in a tris-glycine-SDS buffer before being transferred onto PVDF membranes. The membranes were blocked with 5% skim milk in rinse buffer (0.1% Tween 20, 10 mM Tris-HCl [pH7.5], 0.8% NaCl, and 1mM EDTA) and was incubated with 1st Ab (1:1,000) at 4 °C overnight, and then incubated with HRP-conjugated secondary Ab (1:10,000) for 60 min at room temperature. The immunoblots were visualized with ECL Prime Western Blotting Detection Reagent (GE Healthcare) and detected using a ChemiDoc Touch Imaging System (Bio-Rad).

### Immunoprecipitation

We cultured 5 × 10^5^ of HEK293FT cells were cultured in a 6-well plate for overnight and then performed transfection with expression vectors using Lipofectamine 2000. The total amount of plasmids was maintained at 1 µg by adding empty plasmids. The cells were harvested and washed with PBS after 24 h of transfection, and then lysed with lysis buffer containing protease inhibitor cocktail. The cell lysates were kept on ice for 30 min and were then centrifuged at 15,000 rpm for 20 min at 4 °C. The supernatants were transferred into 1.5 ml tubes. The lysates were pre-treated with protein G sepharose beads at 4°C for 60 min with rotation and then centrifuged to deplete the protein G sepharose beads. Anti-FLAG (1:150) Ab was added to the cell lysates, and then the lysate was incubated for 2 hours at 4 °C with rotation. Washed protein G sepharose beads were added into the lysates containing the Ab, and incubated overnight with rotation. Then, the protein G sepharose beads were collected by centrifugation and washed three times with lysis buffer. The precipitated samples were analyzed using western blots.

### PLA

HEK293 and A549 cells seeded on a glass-bottom plate were fixed with 4% PFA for 15 min at room temperature and permeabilized with 0.3% Triton-X100 in PBS for 60 min at room temperature. Subsequently, the PLA signals were detected by a Duolink In Situ PLA Kit (Sigma-Aldrich), according to the manufacturer’s instructions. Briefly, the cells were blocked with blocking buffer and were labeled with primary Ab and incubated with ligation solution to hybridize the positive and negative probes. The cells were incubated with green or red PLA detection reagent to initiate DNA synthesis with dye-conjugated nucleotides. The indicated number of cells were randomly chosen and observed under confocal microscopy (FV1200: Olympus) to determine the percentage of PLA-positive cells and the number of PLA signals in on cell.

### Reporter gene assay

We seeded 1 × 10^5^ of HEK293 cells into 24-well plates in triplicate and transfected the cells with an expression vector together with p-125 Luc reporter (100 ng/well) and phRL-TK (10 ng/well, Promega) plasmids. The phRL-TK (HSV-thymidine kinase promoter) plasmid encodes *Renilla* luciferase and was used as an internal control. An empty plasmid was added to ensure that each transfection received the same amount of total DNA. After 24 h of transfection, luciferase assays were performed using a Dual Luciferase Assay Kit (Progemga). Luciferase activity was normalized to that of *Renilla* luciferase.

### Quantitative real-time PCR

We seeded 1 × 10^5^ of HEK293 cells into 24-well plates. Then, the cells were transfected with 100 ng of short poly I:C using Lipofectamine 2000 to determine cytokine expression after stimulation. Total RNA was isolated form the cells using TRIzol reagent (Invitrogen), according to the manufacture’s protocol. cDNA was generated from the total RNA using a High-Capacity cDNA Reverse Transcription Kit (Applined Biosystems). The target mRNA was quantified with Power SYBR Green Master Mix (Applied Biosystems) using the ViiA-7 Real-Time PCR system (Applied Biosystems), and the expression levels were normalized to GAPDH.

### Lentivirus production

HEK293FT cells (2 × 10^6^) were seeded on a 10 cm dish and incubated at 37°C for 1 day. GFP, LGP2 and LGP2 4KR ORFs were cloned into a FUIPW lentiviral vector and co-transfected with packaging plasmids into HEK293FT cells. After 3 days of transfection, the viruses were harvested and filtered (0.22 µm filter, Millipore) and used to infect HEK293 cells in the presence of polybrene (10 µg/ml). The infected cells were selected with the application of puromycine (1 µg/ml) for 3 days. Human ACE2 cDNA clone was synthesizes using HepG2 total RNA. The expression of ACE2 in HEK293 ACE2 cells was confirmed by RT-qPCR.

### Nano-liquid chromatography-tandem mass spectrometry (LC-MS/MS)

FLAG-tagged LGP2 and Riplet expression vectors were transfected into HEK293 cells. After 24 h of transfection, the cell lysate was prepared, and LGP2 was purified using anti-FLAG monoclonal Abs. The isolated LGP2 was subjected to SDS-PAGE and stained with CBB. LGP2 bands were excised from the gel. Then, the proteins in each gel slice were subjected to reduction with 10 mM dithiothreitol (DDT), at 56 °C for 1 h, alkylation with 55 mM iodoacetamide at room temperature for 45 min in the dark, and digestion with 10 µg/ml modified trypsin (Promega) at 37°C for 16 h. The resulting peptides were extracted with 1% trifluoroacetic acid and 50% acetonitrile, dried under a vacuum, and dissolved in 2% acetonitrile and 0.1% formic acid. The peptides were then fractionated by C18 reverse-phase chromatography (Advance Nanoflow UHPLC System; AMR Inc.) and applied directly into a hybrid linear ion trap mass spectrometer (LTQ Orbitrap Velos Pro; Thermo Fisher Scientific) with Advanced Captive Spray SOURCE (AMR Inc.) as the ion source. The mass spectrometer was programmed to perform 11 successive scans, with the first consisting of a full MS scan from 350–1,800 m/z by FT-ICR at a resolution of 60,000. Scans 2–11 were of data-dependent scans of the top ten abundant ions obtained in the first scan, by ion trap. Automatic MS/MS spectra were obtained from the highest peak in each scan by setting the relative collision energy to 35% and the exclusion time to 20 s for molecules in the same m/z value range. The molecular masses of the resulting peptides were searched against the Uniprot Proteome Homo sapiens database (downloaded 2018.12.13) using the Mascot v2.6 program via Proteome Discoverer v2.2 (Themo Fisher Scientific) with the false discovery rate (FDR) set at 0.01. Cysteine carbamidomethylation was set as a fixed modification. The oxidation of methionine, acetylation of protein N-termini, and ubiquitination of lysine were set as variable modifications. The number of missed cleavage sites was set as 2.

### Quantification and statistical analysis

All qPCR assays and reporter gene assays were performed in triplicate (n = 3). Error bars represent SD. Statistical significance (p-value) was determined using a two-tailed Student’s *t*-test, one-way ANOVA, or two-way ANOVA in Prism v7.0a (GraphPad Software) and MS-Excel (Microsoft Corp) software. *p < 0.05.

